# Degradome analysis to identify direct protein substrates of small-molecule degraders

**DOI:** 10.1101/2024.01.28.577572

**Authors:** Marco Jochem, Anna Schrempf, Lina-Marie Wagner, Jose Cisneros, Amanda Ng, Georg E. Winter, Jeroen Krijgsveld

**Affiliations:** German Cancer Research Center (DKFZ), Heidelberg, Germany; CeMM Research Center for Molecular Medicine, Vienna, Austria; Heidelberg University, Faculty of Medicine, Heidelberg, Germany

## Abstract

Targeted protein degradation (TPD) has emerged as a powerful strategy to selectively eliminate cellular proteins using small-molecule degraders, offering therapeutic promise for targeting proteins that are otherwise undruggable. However, a remaining challenge is to unambiguously identify primary TPD targets that are distinct from secondary downstream effects in the proteome. Here we introduce an approach that combines stable isotope labeling and click-chemistry for selective quantification of protein degradation by mass spectrometry, excluding confounding effects of altered transcription and translation induced by target depletion. We show that the approach efficiently operates at the time scale of TPD (hours) and we demonstrate its utility by analyzing the Cyclin K degraders dCeMM2 and dCeMM4, which induce widespread transcriptional downregulation, and the GSPT1 degrader CC-885, an inhibitor of protein translation. Additionally, we apply it to characterize compound 1, a previously uncharacterized degrader, and identify the zinc-finger protein FIZ1 as a degraded target.

## Introduction

Small-molecule degraders are a category of drugs that exploit the ubiquitin machinery by recruiting E3 ligases to target proteins of interest for their ubiquitination and subsequent degradation by the proteasome (Békés et al., 2022; Chamberlain & Hamann, 2019; Zhao et al., 2022). This mechanism is very different from traditional drugs that bind and inhibit their targets at functional sites, and therefore TPD is a promising therapeutic strategy particularly in cases where achieving specificity and druggability has proven challenging. In addition, small-molecule degraders offer advantageous pharmacodynamics, benefiting from their catalytic mode of action that obviates continued target engagement. Moreover, substrate degradation neutralizes both the enzymatic and scaffolding functions of their targets and might hence offer a more profound perturbation. Clinically approved examples of small-molecule degraders are found within the immunomodulatory imide drug (IMiD) family, including thalidomide and its analogues lenalidomide and pomalidomide (Ito et al., 2010; Krönke et al., 2014; Lu et al., 2014). IMiDs bind to the E3 ligase CRBN and thereby recruit the zinc finger transcription factors IKZF1 and IKZF3, causing their ubiquitination and ensuing degradation by the proteasome. Of note, IMiDs have been successfully used to treat blood cancers such as multiple myeloma. Additionally, numerous small-molecule degraders currently undergo clinical trials, targeting a wide range of proteins, such as Androgen and Oestrogen receptor, BCL-XL, IRAK4, STAT3, BTK, TRK, BRD9, IKZF1/2/3, and GSPT1 (Chirnomas et al., 2023; Mullard, 2021).

Based on their chemical structure and mechanism of action, small-molecule degraders can be divided into two major classes: proteolysis targeting chimeras (PROTACs) and molecular glues (Hanzl & Winter, 2020). PROTACs are bifunctional molecules composed of two warheads that selectively bind to the E3 ligase and the protein of interest, respectively, connected by a flexible linker. This approach offers a high degree of synthetic flexibility due to the large number of different possible combinations of warheads. In contrast, molecular glues enhance direct protein-protein interactions between the E3 ligase and the target protein to either boost the ubiquitination of existing substrates or enable the ubiquitination and degradation of neo-substrates (Geiger et al., 2022; Gerry & Schreiber, 2020). Molecular glues are typically smaller in size than PROTACs, making them more amenable to pharmacological administration, however their rational design is complicated by the fact that target engagement is difficult to predict, usually necessitating complex screening strategies to identify novel compounds (Hanzl et al., 2023; Mayor-Ruiz et al., 2020; Ng et al., 2023).

Since small-molecule degraders act at the protein level, proteomic methods have become essential to identify their targets and to determine target specificity. Liquid chromatography mass spectrometry (LC-MS) based methods employing label-free or tandem mass tag (TMT)-based multiplexing strategies are routinely used to identify drug-induced proteomic changes without requiring prior knowledge of the drug’s targets (Meissner et al., 2022; Sathe & Sapkota, 2023; Scholes et al., 2021). However, in the context of TPD, conventional global proteomics approaches face the challenge of distinguishing proteins that are degraded as direct drug targets from those that are downregulated due to indirect downstream effects, illustrated by the observation that many drugs alter overall proteome composition (Mitchell et al., 2023). This can be expected especially when targeting transcriptional or translational regulators, which complicates the identification of direct drug targets and inflates the number of candidates that need to be validated independently, as we noted in our own work (Mayor-Ruiz et al., 2020).

To overcome these challenges, we introduce a proteomics workflow tailored to selectively analyze drug-induced protein degradation while excluding proteomic alterations that arise from indirect transcriptional and translational effects. We apply this degradome strategy to characterize direct targets of established and uncharacterized protein degraders.

## Results

To identify direct targets of TPD, we developed a strategy to specifically monitor protein degradation at proteomic scale (i.e., for ‘degradomic’) that we adapted from our previous method to investigate newly synthesized proteins (Eichelbaum et al., 2012, Figure 1a). Exploiting stable isotope labeling by amino acids in cell culture (SILAC), one key step involves exposing SILAC-intermediate-labeled cells to both azidohomoalanine (AHA, a clickable methionine analogue) and either SILAC-light or SILAC-heavy amino acids for an 8-hour pulse period. This creates a pool of labeled proteins across the proteome whose abundance is monitored after switching cells back to SILAC-intermediate media, while simultaneously adding a protein degrader or vehicle-treatment (to SILAC-heavy and SILAC-light labelled cells, respectively). Subsequently, AHA-containing proteins from the labeled pool are enriched using click-chemistry and identified and quantified by mass spectrometry. Crucially, proteins that are newly synthesized after drug addition do not incorporate AHA and differ in their SILAC label status (intermediate). Hence, these proteins are not enriched and can otherwise be distinguished from SILAC-heavy and SILAC-light proteins originating from the labeled protein pool. Therefore this design allows for the selective study of drug-induced protein degradation that excludes secondary transcriptional and translational effects that may occur after drug administration, as a consequence of target (or off-target) degradation. We reasoned that this degradomics approach should be ideally placed to identify direct targets of TPD.

**Figure 1:**
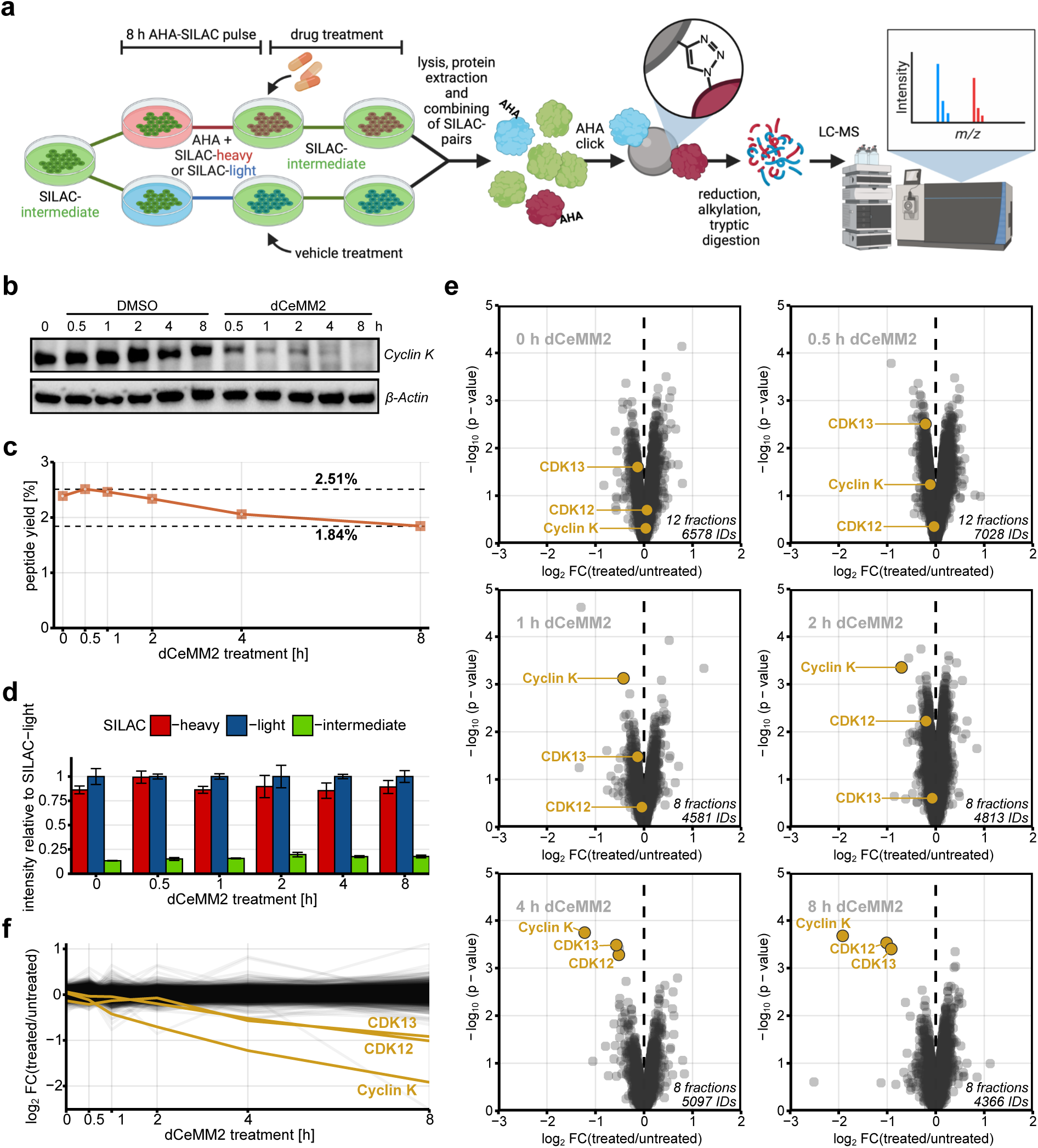
Degradome proteomics. **(a)** Workflow scheme: Fully SILAC-intermediate labeled cells are pulse-labeled with both AHA and SILAC-heavy or SILAC-light amino acids for 8 h. The media is changed back to SILAC-intermediate, and the former heavy-channel is treated with the drug of interest, while the former light-channel is mock-treated. Cells are lysed, proteins extracted, and the corresponding SILAC-channels combined. The pulsed proteins are enriched for AHA via click-chemistry, reduced, alkylated, and digested by trypsin. The resulting peptides are analyzed by LC-MS/MS for protein identification and quantification to determine drug-induced protein degradation inferred from heavy-over-light SILAC-ratios. **(b)** Degradation of Cyclin K in response to dCeMM2 determined by immunoblot analysis. KBM7 cells were either treated with 2.5 µM dCeMM2 or with DMSO for the indicated time intervals. Upper panel: anti-Cyclin K antibody. Lower panel: anti-β-Actin (loading control). **(c)** Peptide yields after enriching 1.2 mg of protein for AHA while performing degradome proteomics. KBM7 cells were treated with 2.5 µM dCeMM2 or DMSO for 0, 0.5, 1, 2, 4 or 8 h. Protein and peptide yields were monitored via BCA-assay. **(d)** Summed peptide intensities of each SILAC-channel relative to the corresponding SILAC-light channel. Error bars represent standard errors of the means. **(e)** Volcano plots showing differential protein stabilities in response to the dCeMM2 treatment time course. Limma statistical analysis was used on MaxQuant-normalized heavy-over-light SILAC-ratios. Highlighted are the three known targets of dCeMM2: Cyclin K, CDK12 and CDK13. **(f)** Line plot visualizing global protein stabilities over the dCeMM2 time course. Note that each line represents one protein and that only proteins quantified in all timepoints are shown.

To test this, we applied our degradome workflow to investigate the small-molecule degrader dCeMM2, which we previously discovered through a screening approach for molecular glue degraders (Mayor-Ruiz et al., 2020), and which targets Cyclin K and its interaction partners, the transcriptional kinases CDK12 and CDK13 (Figure 1b). Given that the degradation of Cyclin K, CDK12, and CDK13 leads to widespread transcriptional downregulation of other genes (Mayor-Ruiz et al., 2020), dCeMM2 provided an ideal test case to evaluate the performance of our approach. We pulse-labelled human KBM7 cells, added the drug, and sampled cells at 6 time points during an 8-hour time course treatment. Upon digestion of AHA-enriched proteins we observed a noticeable decrease in peptide yield with prolonged drug treatment (Figure 1c), which is expected since no newly synthesized proteins can incorporate AHA, yet all proteins in the AHA-labelled pool progressively undergo both basal and drug-induced degradation. As anticipated, SILAC-intermediate labelled peptides represented by far the smallest fraction of all peptides after enrichment (Figure 1d), demonstrating the efficacy of the workflow in excluding proteins synthesized before and after the 8-hour SILAC-AHA pulse period. dCeMM2-induced proteome effects were quantified from SILAC-heavy/SILAC-light ratios, revealing that degradation of the intended target Cyclin K occurred already after 1 hour of drug treatment, followed by the degradation of CDK12 and CDK13 after 4 hours (Figure 1e, Supplementary Data 1). In fact, these three targets of dCeMM2 were the only downregulated proteins throughout the treatment time course among 4000-7000 identified proteins, indicating the ability of the workflow to selectively identify degradation targets of small molecule degraders (Figure 1f).

To compare the performance of our workflow with commonly used global proteomic methods, we performed a degradome proteomics experiment in which cells were treated with dCeMM2 and dCeMM4 for 12-hours and compared the results with data previously obtained using a TMT-based approach under otherwise identical conditions (Mayor-Ruiz et al., 2020, Figure 2a and b, Figure S1a, Supplementary Data 2). While several proteins were downregulated in the TMT-dataset along with Cyclin K, CDK12 and CDK13, virtually no off-targets were observed in the corresponding degradome dataset. In fact, using our degradome workflow, we found that the three primary targets of dCeMM2 and dCeMM4 almost exclusively ranked at the very top of degraded proteins, while the false-positives in the TMT-approach tended to be stabilized rather than degraded (indicated in red in Figure 2a and b). A notable exception was the mRNA export adaptor FYTTD1/UIF, which both proteomics approaches detected as downregulated in response to both drugs, signifying it as another target of degradation. Interestingly, FYTTD1 was previously determined to be a Nedd8-dependent substrate of Cullin-RING ubiquitin ligases (Liao et al., 2011), and it was identified as an off-target of both the BET-degrader JQ-1 (leading to downstream arrest in protein synthesis, Savitski et al., 2018), and of a CDK9 degrader acting via the E3 ligase KEAP1 (Pei et al., 2023).

**Figure 2:**
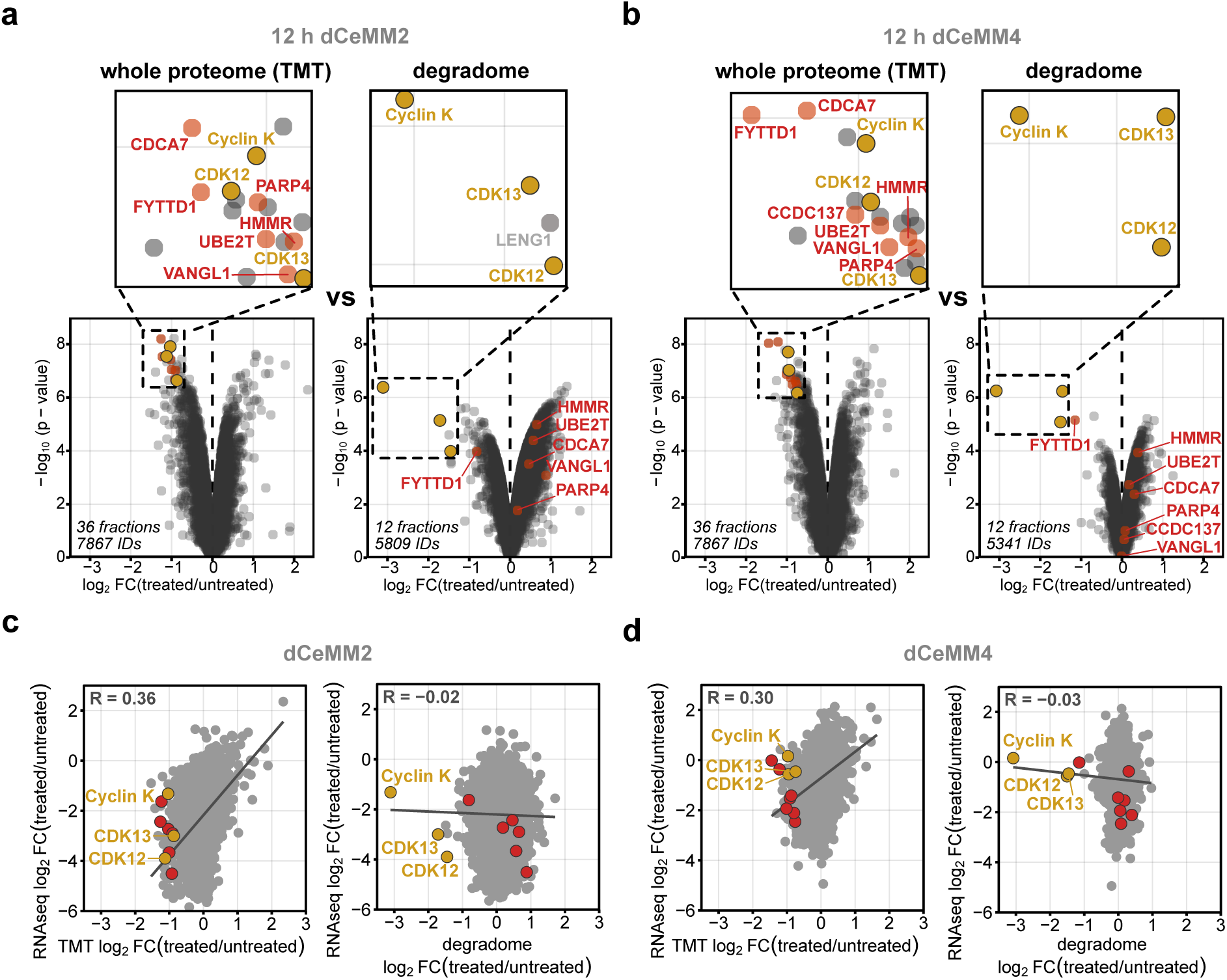
Benchmarking degradome proteomics. **(a, b)** Comparing degradome proteomics (right) with TMT-based whole proteome analysis (left) for Cyclin K degraders. KBM7 cells were treated with dCeMM2 (2.5 µM) or dCeMM4 (3.5 µM) for 12 h in both approaches. Whole proteome data were taken from Mayor-Ruiz et al., 2020. Highlighted in yellow are the three described degradation targets of dCeMM2 and dCeMM4: Cyclin K, CDK12 and CDK13. The margins of the zoom windows include all proteins with a more extreme fold-change and a better p-value than Cyclin K, CDK12 and CDK13. Highlighted in red are the proteins that localized inside the zoom window of the whole proteome analysis and that were identified using degradome proteomics. **(c, d)** Pearson correlation between the transcriptome (Mayor-Ruiz et al., 2020) and the whole proteome or the degradome of dCeMM2- and dCeMM4-treated cells. The treatment time for the whole proteome and the degradome analysis was 12 h. The treatment time for the transcriptome analysis was 5 h.

Many of the proteins that exhibited significant downregulation in the whole proteome dataset were also downregulated at the transcript level (Figure S1b and c, RNAseq data by Mayor-Ruiz et al., 2020). We even observed a positive correlation between the transcriptome and whole proteome for both dCeMM2 and dCeMM4-treated cells (R = 0.36 and R = 0.3), indicating that a substantial portion of drug-induced proteome changes was driven by transcriptional regulation (Figure 2c and d). In contrast, the degradome dataset showed no correlation with the transcriptome data (R = - 0.02 and R = -0.03), underscoring that our approach is not confounded by such secondary effects.

We next turned to CC-885, a pleiotropic molecular glue degrader from the IMiD family that targets the small GTPase GSPT1 (Matyskiela et al., 2016). Since GSPT1 (also called eRF3 for eukaryotic release factor 3) functions by interacting with the translation termination factor eRF1 to mediate protein translation, CC-885 treatment leads to extensive remodelling of the proteome, masking GSPT1 among multiple other downregulated proteins (Powell et al., 2020). Reflecting this challenge to distinguish direct from indirect targets, in fact GSPT1 was originally identified as a substrate of CC-885 by affinity pulldown of tagged CRBN, not by global proteomics (Matyskiela et al., 2016). To evaluate whether our degradome approach could lead to a more unambiguous identification of direct CC-885 targets by negating secondary effects caused by GSPT1 depletion, we conducted side-by-side analyses of different proteomic methods on CC-885-treated cells. Analysing the whole proteome via label-free data independent acquisition (DIA)-MS, we observed the downregulation of numerous proteins, including the anticipated targets GSPT1, IKZF1 and IKZF3 (Figure 3a, Supplementary Data 3). This sharply contrasted with the degradome analysis, where GSPT1, IKZF1, and IKZF3 stood out as the prime targets, while the other proteins that were downregulated in the global proteome were not affected (Figure 3b, Supplementary Data 3). Interestingly, a nascent proteome analysis that selectively labels and enriches for newly synthesized proteins (for experimental scheme see Figure S2), showed that CC-885 treatment induced a profound and global downregulation of proteins, including those observed in the global proteome, as well as GSPT1, IKZF1, and IKZF3 (Figure 3c, Supplementary Data 3), consistent with a generic role of GSPT1 in translation termination. Additionally, the degradome and the nascent proteome approaches identified the downregulation of the GSPT1-homologue GSPT2, which has been previously found as a target of CC-885 (Powell et al., 2020), and the E3 ubiquitin ligase RNF166, observed as an off-target of several IMiDs (Donovan et al., 2020; Kozicka and Thomä, 2021). Collectively, these data show the ability of our degradome proteomics approach to identify direct substrates of TPD even if they have a profound role in regulating proteome homeostasis, such as GSPT1.

**Figure 3:**
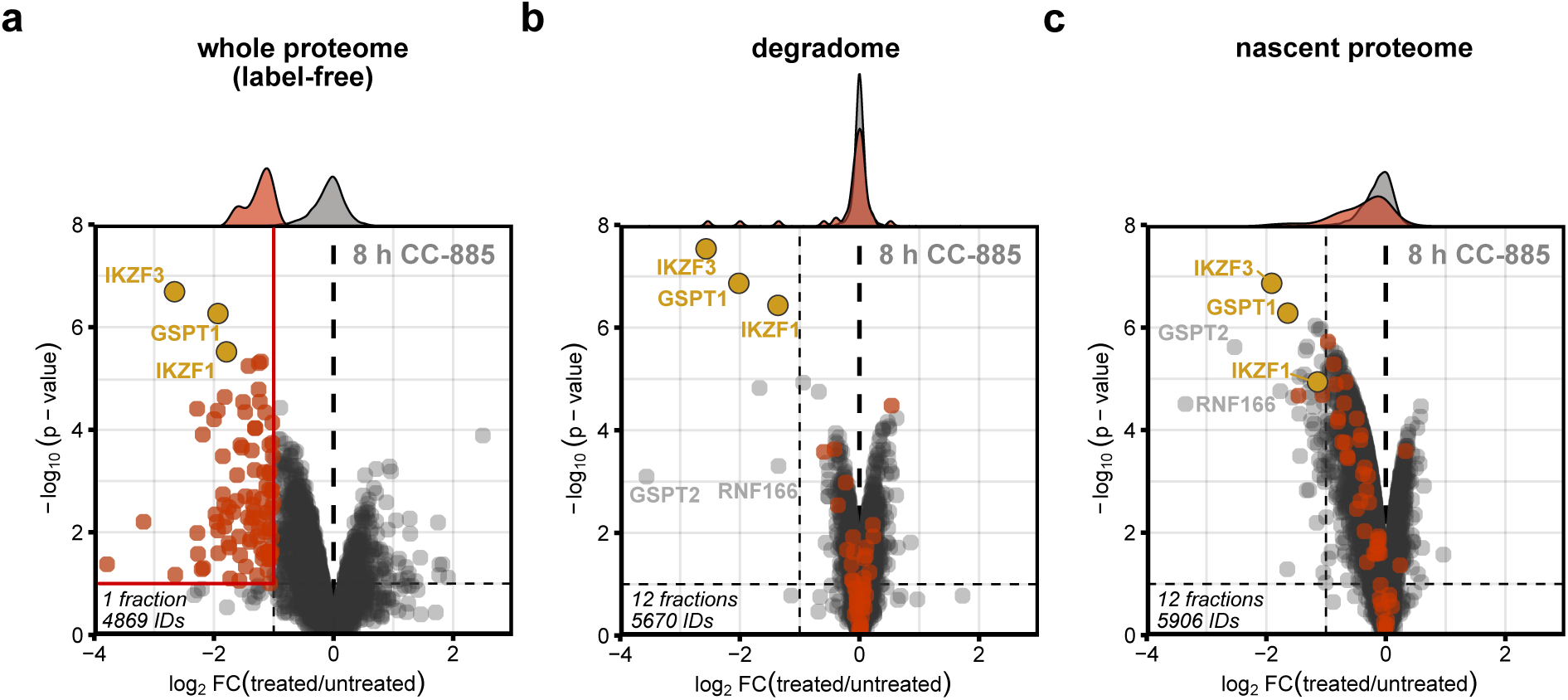
Proteome response to CC-885 treatment. **(a)** Label-free whole proteome, **(b)** degradome, and **(c)** nascent proteome analysis for GSPT1 degrader CC-885. Jurkat cells were treated with 100 nM CC-885 for 8 h in all three approaches. Highlighted in yellow are three known targets of CC-885, and in red all proteins that were downregulated (log_2_FC(treated/untreated)<-1 and -log_10_(p-value)>1) in the whole proteome dataset. Density plots visualize the fold-change distribution of all proteins (in grey) and of the yellow- and red-labelled proteins (in red).

Next, we used degradome proteomics to characterise the previously disclosed IMiD-based degrader compound 1, yet whose targets are unknown (Figure 4a, Bradner et al., 2017). Upon treatment of Jurkat cells with compound 1, two common IMiD targets IKZF1 and IKZF3 were among the most significantly degraded proteins (Figure 4b, Supplementary Data 4), as confirmed by western blot (Figure 4c). ZFP91 was at the border of significance (Figure 4b) and is also a common target of IMiDs (Kozicka and Thomä, 2021), collectively indicating that we accessed the expected target space. In addition, we observed the degradation of FIZ1 (Flt3-interacting zinc finger protein 1), which, like many other IMiD targets, is a zinc finger-containing protein, but which to our knowledge has not been previously described as a substrate of other small-molecule degraders. FIZ1 plays a role in FMS-like receptor tyrosine kinase 3 (FLT3) signalling (Tsuruyama et al., 2010; Wolf & Rohrschneider, 1999), functions as a transcriptional repressor (Mali et al., 2007, 2008; Mitton et al., 2003), and is associated with pro-proliferative effects in keratinocytes (Larivière et al., 2014). We confirmed the compound 1-induced degradation of FIZ1 through immunoblot analysis in both HEK293T and Jurkat cells (Figure 4d and e). Furthermore, we demonstrated via luminescence assay that the rate of degradation is dose-dependent (Figure 4f). In line with a CRBN-dependent mechanism of action, we observed that the compound 1-induced degradation of FIZ1 was impeded when inhibiting the proteasome, ubiquitination, or neddylation, (Figure 4g). Collectively, this establishes FIZ1 as a genuine target of compound 1. Investigating an extended panel of IMiDs, we found that both pomalidomide and CC-90009 are also capable of degrading FIZ1 (Figure 4h). In conclusion, degradome analysis efficiently identifies direct targets of small-molecule degraders by negating secondary proteome effects induced by target depletion, thereby filling an important need for target identification of novel protein degraders.

**Figure 4.**
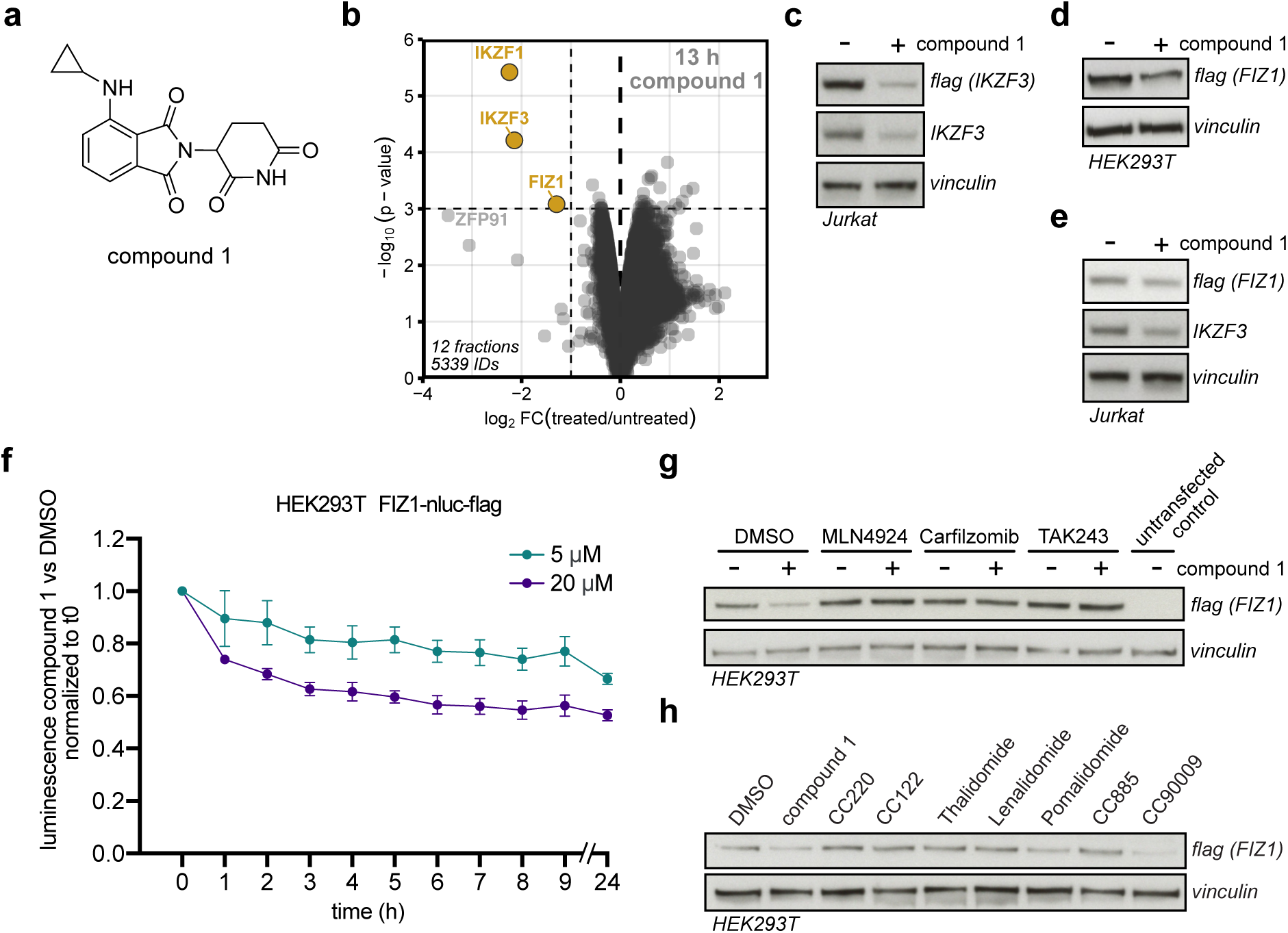
Validation of FIZ1 as a degradation target of compound 1. **(a)** Chemical structure of small molecule degrader compound 1. **(b)** Degradome proteomics under compound 1 treatment. Jurkat cells were treated for 13 h with 5 µM compound 1 or DMSO. Limma statistical analysis was used on MaxQuant-normalized heavy-over-light SILAC-ratios. Downregulated proteins are highlighted in yellow (log_2_FC(treated/untreated) < -1 and -log_10_(p-value) > 3). **(c)** Degradation of IKZF3-nluc-flag and endogenous IKZF3 in response to compound 1 treatment determined by immunoblot analysis. Jurkat cells were treated with 10 µM compound 1 or DMSO for 15 h. Antibodies used: anti-flag, anti-IKZF3 and anti-vinculin (loading control). **(d)** Degradation of FIZ1-nluc-flag in response to compound 1 treatment determined by immunoblot analysis in Hek293T cells and **(e)** in Jurkat cells. Cells were treated with 10 µM compound 1 or DMSO for 16 h. Antibodies used: anti-flag, anti-IKZF3 (positive control) and anti-vinculin (loading control). **(f)** Luminescence assay in HEK293T cells treated with either compound 1 (5 µM, 20 µM) or DMSO for the indicated times. **(g)** Degradation of FIZ1-nluc-flag in response to 20 µM compound 1 treatment for 13 h in HEK293T cells under inhibition of neddylation (1 µM MLN4924 = NAE inhibitor), the proteasome (1 µM Carfilzomib) or ubiquitination (10 µM TAK243 = UBA1 inhibitor), determined by immunoblot analysis. Upper panel: anti-flag antibody. Lower panel: anti-vinculin (loading control). **(h)** Degradation of FIZ1-nluc-flag in response to different small molecule degraders determined by immunoblot analysis. HEK293T cells were treated with DMSO, 20 µM compound 1, 10 µM CC220, 10 µM CC122, 10 µM Thalidomide, 10 µM Lenalidomide, 10 µM Pomalidomide, 10 µM CC-885 or 10 µM CC-90009 for 15 h.

## Discussion

In this study, we have introduced and validated a proteomics-based workflow designed to selectively identify targets of small-molecule protein degraders. A key feature of our approach is the incorporation of a SILAC and AHA pulse period prior to drug treatment, which creates an isolated protein pool that is not influenced by altered protein translation resulting from subsequent depletion of TPD targets. Unlike conventional whole proteome analyses and pulse-chase experiments to monitor protein decay under steady-state conditions (McShane et al., 2016), our method is purpose-built to discern acute perturbation-induced changes in protein stability, and it is therefore tailored to the target-discovery of protein degraders. We showed its efficacy by identifying targets of known degraders (CeMM2, CeMM4, CC-885), even if their targets themselves impact proteome homeostasis (Cyclin K, GSPT1, Figure 2a and b, Figure 3). Although analysis of nascent proteins has been proposed as a method to identify TPD targets (An et al., 2017), we demonstrated that direct analysis of protein degradation led to more unambiguous target identification while negating secondary effects in the proteome (Figure 3b and c). Moreover, we establish that degradome analysis operates at the rapid time scale of TPD (hours, Figure 1e), offering the possibility to limit duration of drug treatment and further minimize the risk of capturing secondary effects. Since GSPT1 degraders are tested in various clinical trials, degradome analysis may be part of such programs to evaluate drug specificity.

We envision that the utility of degradome proteomics extends to other areas of TPD research, or even more generally to the study of protein stability in various other biological contexts. For instance, the mechanisms and cargo of lytic organelles may be characterized using autophagy targeting chimeras (AUTACs, Takahashi et al., 2019) and lysosome targeting chimeras (LYTACs, Banik et al., 2020). Conversely, the recruitment of deubiquitinases (DUBs) to other proteins through DUB-targeting chimeras (DUBTACs) can lead to their targeted stabilization (TPS), another novel therapeutic concept that is especially relevant in pathologies that are driven by excessive protein degradation (Henning et al., 2022). Although not specifically assessed in this study, we anticipate that our method has the capacity to measure drug-induced protein stabilization as well. Our method also has the potential to assist in the identification of endogenous substrates of E3 ligases, which is important for drug development but has proven challenging (Iconomou & Saunders, 2016).

Currently, a bottleneck of our method resides in the use of offline peptide fractionation, a time-intensive and laborious process that was used here to achieve comprehensive proteome coverage in data-dependent acquisition (DDA). However, we anticipate that this step can be omitted with the emergence of data-independent acquisition (DIA) optimized for SILAC-labelled proteins (Pino et al., 2021). In fact, we have recently implemented such an approach in combination with a semi-automated workflow to perform AHA-based protein enrichment for the investigation of perturbation-induced effects on protein translation (Borteçen et al., 2023). The prospect of automated degradome analyses presents an exciting opportunity to develop a powerful tool tailored for high-throughput drug screening in TPD research.

## Methods

### Small-molecule degraders

MLN4924 (THP Medical #HY-70062), Carfilzomib (Selleck Chemicals #S2852), TAK243 (MedChem Express #HY-100487), CC220 (Selleck Chemicals #S8760), CC122 (Selleck Chemicals #S7892), Thalidomide (Sigma-Aldrich #T144), Lenalidomide (Sigma-Aldirch #SML2283), Pomalidomide (THP Medical Product #HY-10984), CC-885 (MedChem Express #HY-101488), CC-90009 (MedKoo #207005).

### Synthesis of compound 1

Reactions were monitored by thin-layer chromatography (TLC) using pre-coated silica gel plates F-254. Proton nuclear magnetic resonance (^1^H NMR) spectra and carbon nuclear magnetic resonance (^13^C NMR) spectra were recorded on Bruker AV III 600 instrument from the NMR Facility of the University of Vienna. The NMR peak multiplicities are denoted as follows: s, singlet; d, doublet; dd, doublet of doblets; m, multiplet. Column chromatography was carried out using Biotage Selekt over Biotage Sfar Silica D column cartridges employing Merck silica gel (Kieselgel 60, 63-200 mm). HRMS (ESI-TOF) analyses were performed on Bruker timsTOF flex at the MS Facility of the University of Vienna.

A mixture of 2-(2,6-dioxopiperidin-3-yl)-4-fluoroisoindoline-l,3-dione (20 mg, 0.0724 mmol, 1.0 eq.), cyclopropylamine (0.080 mmol, 1.1 eq.) and DIPEA (0.141 mmol, 1.9 eq.) in N-methylpyrrolidinone (0.1 M) was heated to 90 0C overnight. The reaction mixture was cooled to room temperature and diluted in ethyl acetate. The organic layer was washed with saturated solution of NaHC03, water and brine, and subsequently dried over anhydrous sodium sulfate and concentrated under reduced pressure. The residue was purified by flash column chromatography on silica gel using dichloromethane/methanol as solvent system to afford the title compound as a yellow powder.

4-(cyclopropylamino)-2-(2,6-dioxopiperidin-3-yl)isoindoline-1,3-dione. 22 mg. Yield 70%. Characterization data correspond to previously reported. ^1^H NMR (600 MHz, DMSO) δ 11.09 (s, 1H), 7.65 (dd, *J* = 8.5, 7.1 Hz, 1H), 7.36 (d, *J* = 8.5 Hz, 1H), 7.16 – 7.07 (m, 1H), 6.64 (d, *J* = 2.0 Hz, 1H), 5.05 (dd, *J* = 12.9, 5.5 Hz, 1H), 2.95 – 2.82 (m, 1H), 2.62 – 2.56 (m, 2H), 2.55 – 2.46 (m, 1H), 2.07 – 1.97 (m, 1H), 0.86 – 0.79 (m, 2H), 0.61 – 0.52 (m, 2H). ^13^C NMR (151 MHz, DMSO) δ 172.8, 170.0, 168.7, 167.3, 146.8, 136.1, 132.0, 118.3, 111.3, 109.7, 48.5, 30.9, 24.1, 22.1, 7.3. HRMS (ESI) [M + H]^+^ calcd for C16H15N3O4, 314.1135, found 314.1136.

### Cell culture, lentivirus production and generation of stable cell lines

All cells were cultured at 37°C in a humidified 5% CO2-atmosphere. Unlabeled KBM7 cells were maintained in Iscove’s Modified Dulbecco’s Medium (IMDM) supplemented with 10% (v/v) FBS and 1x Pen-Strep (Gibco #15140-122). Unlabeled Jurkat cells were maintained in Roswell Park Memorial Institute (RPMI) 1640 medium supplemented with 10% (v/v) FBS and 1x Pen-Strep. HEK293 cells (a gift by the Bradner Lab) were maintained in Dulbecco’s Modified Eagle Medium (DMEM) supplemented with 10% (v/v) FBS and 1x Pen-Strep. SILAC-intermediate labeled KBM7 and Jurkat cells were maintained in RPMI 1640 for SILAC (Thermo Scientific #A33823) supplemented with 10% (v/v) dialyzed FBS (dFBS, Gibco #26400-044), 1x Pen-Strep, 200 mg/l L-proline, 1x GlutaMAX (Gibco #35050061), 200 mg/l SILAC-intermediate L-lysine (Silantes #211103913) and 40 mg/l SILAC-intermediate L-arginine (Silantes #201203902). For lentiviral production, HEK293T cells were seeded in 6-well plates and transfected at approx. 80 % confluency with 0.69 µg target vector, 0.17 µg VSV.G (Addgene #14888) and 0.34 µg psPAX2 (Addgene #12260) using PEI following standard protocol. 72 h later, supernatant containing virus was harvested, centrifuged at 2000 rpm for 5 min to remove cell debris. Lentivirus was then aliquoted and stored at – 80 °C until transduction of 0.25 × 10^6 cells in 1 ml of media plus virus in 24 well plates with the addition of 8 µg/ml polybrene (Sigma) and spin inoculation for 1 h at 2,000 r.p.m. Antibiotic selection was performed 24 h after transduction with 1 µg/ml puromycin.

### Plasmids and cloning

For overexpression of IKZF3 and FIZ1, full-length IKZF3 and FIZ1 was synthesized by Genscript and cloned into a modified pLEX305 backbone harboring a nluc-3x-FLAG tag via Gateway cloning (Thermo Fisher Scientific) following manufacturer’s recommendations.

### Western blot

Cells were harvested into ice-cold PBS, pelleted, and resuspended in RIPA lysis buffer containing 50 mM Tris-HCl pH 7.5, 1% (v/v) NP-40, 0.25% (w/v) Na-Deoxycholate, 0.1% (w/v) SDS, 150 mM NaCl, 1 mM EDTA, 10% (v/v) glycerol and 1× Halt or 1x cOmplete^TM^ EDTA-free protease inhibitor cocktail (Roche #11873580001). Lysates were sonicated with a probe sonicator and cleared via centrifugation at 15,000x g for 5 min at 4°C. Protein concentrations were measured with either the Pierce^TM^ BCA Protein Assay Kit (Thermo Scientific #23225) or the Pierce^TM^ Rapid Gold BCA Protein Assay Kit (Thermo Scientific #A53225PR). One part of cleared lysate was combined with three parts of 4x Laemmli Sample Buffer (Bio-Rad #1610747) containing 400 mM DTT. Samples were heated to 90°C for 10 min, vortexed and cooled down to room temperature. Equal protein amounts were loaded onto 4-15% Mini-PROTEAN^®^ TGX^TM^ Precast Protein Gels (Bio-Rad #4561086DC) submerged in 1x TGS-Buffer (Bio-Rad #1610772). Precision Plus Protein^TM^ Dual Color Standard (Bio-Rad #4561084DC) was loaded as a molecular weight reference and the electrophoresis performed until the dye reached the end of the gel. Proteins were transferred onto an Amersham Hybond^TM^ 0.45 µm PVDF Blotting Membrane (Cytiva #15259894). Standard Western Blot analysis was performed using primary antibodies against Cyclin K (1:5000, Bethyl Laboratories #A301-939A), β-Actin (1:4000, Cell Signaling #4970), α-Tubulin (1:2000, GeneTex #GT114), Vinculin (1:5000, Santa Cruz #sc-6954), Flag (DYKDDDDK, 1:1000, CellSignaling #2368), IKZF3 (1:1000, Novus Biologicals #NBP2-24495), CRBN (1:2000, kind gift of R. Eichner and F. Bassermann). Anti-rabbit IgG-HRP (1:10000, Santa Cruz Biotechnology #sc-2357 or Jackson ImmunoResearch #111-035-003) and anti-rabbit IgG-HRP (1:10000, Santa Cruz Biotechnology #sc-516102 or Jackson ImmunoResearch #111-035-003) were used for enhanced chemiluminescence detection with either Clarity Western ECL Substrate (Bio-Rad #1705061) or Clarity Max Western ECL Substrate (Bio-Rad #1705062).

### Pulsed-SILAC-AHA labeling for degradome analysis

Fully SILAC-intermediate labeled KBM7 or Jurkat cells were seeded into RPMI SILAC-intermediate medium and grown overnight. The next day, the old media was removed and the cells were washed with warm PBS two times. Cells were deprived of methionine by incubating them for 1 h in methionine-free depletion medium consisting of SILAC-RPMI 1640 (AthenaES #AES-0432) supplemented with 10% (v/v) dFBS, 10 mM HEPES, 1 mM sodium pyruvate (Gibco #11360070), 300 mg/l L-proline, 50 mg/l L-leucine and 1x GlutaMAX (Gibco #35050061). After removal of the depletion medium, half of the cells were pulse-labelled in AHA-SILAC-heavy medium consisting of depletion media additionally supplemented with 200 mg/l SILAC-heavy L-lysine (Silantes #211604102), 40 mg/l SILAC-heavy L-arginine (Silantes #201604102) and 0.1 mM L-azidohomoalanine (Click Chemistry Tools #1066-1000). The other half of the cells were labelled in AHA-SILAC-light medium containing regular L-lysine, regular L-arginine and L-azidohomoalanine at the same concentrations as the heavy media. After 8 h of AHA-SILAC-pulse labeling, cells were transferred into RPMI SILAC-intermediate medium again and divided into multiple smaller cultures (one 10 cm dish per timepoint, replicate and SILAC-channel). The heavy-pulsed cells were treated with the drug of interest for the indicated amount of time, while the light-pulsed cells were treated with DMSO. Cells were harvested by centrifugation, washed with ice-cold PBS and pellets stored at -20°C until lysis.

### Pulsed SILAC-AHA labeling for translatome analysis

Fully SILAC-intermediate labeled Jurkat cells were seeded into RPMI SILAC-intermediate medium and grown overnight. The next day, the old medium was removed and the cells were washed with warm PBS two times. Cells were deprived of methionine by incubating them for 1 h in methionine-free depletion medium consisting of SILAC-RPMI 1640 (AthenaES #AES-0432) supplemented with 10% (v/v) dFBS, 10 mM HEPES, 1 mM sodium pyruvate (Gibco #11360070), 300 mg/l L-proline, 50 mg/l L-leucine and 1x GlutaMAX (Gibco #35050061). After removal of the depletion medium, half of the cells were subjected to AHA-SILAC-heavy medium consisting of depletion media additionally supplemented with 200 mg/l SILAC-heavy L-lysine (Silantes #211604102), 40 mg/l SILAC-heavy L-arginine (Silantes #201604102) and 0.1 mM L-azidohomoalanine (Click Chemistry Tools #1066-1000), while the other half of the cells were subjected to AHA-SILAC-light medium containing regular L-lysine, regular L-arginine and L-azidohomoalanine at the same concentrations as the heavy media. Cells were divided into multiple smaller cultures (one 10 cm dish per timepoint, replicate and SILAC-channel) and the heavy-pulsed cells were treated with the drug of interest for the indicated amount of time, while the light-pulsed cells were treated with DMSO. Cells were harvested by centrifugation, washed with ice-cold PBS and pellets stored at -20°C until lysis.

### Click-iT protein enrichment, reduction, alkylation and digestion

Cell pellets from pulsed-SILAC-AHA experiments were resuspended in urea lysis buffer consisting of 200 mM HEPES pH 8, 0.5 M NaCl, 4% (w/v) CHAPS, 8 M Urea and 1x cOmplete^TM^ EDTA-free protease inhibitor (Roche #11873580001). Lysates were sonicated with a probe sonicator and cleared via centrifugation at 15,000x g for 30 min. Protein concentrations were measured with either the Pierce^TM^ BCA Protein Assay Kit (Thermo Scientific #23225) or the Pierce^TM^ Rapid Gold BCA Protein Assay Kit (Thermo Scientific #A53225PR). Equal protein amounts of corresponding SILAC-pairs were combined to a total of 3 mg. AHA-labeled proteins were enriched with the Click-iT Protein Enrichment Kit (Invitrogen #C10416). For this, 100-120 µl of resin were washed with water two times in order to remove the storage solution. The beads were resuspended in 100 µl water, combined with the lysate and the volume adjusted to 1.9 ml with urea lysis buffer. To start the Click-iT reaction, 10 µl of 200 mM Cu(II)SO_4_, 62.5 µl of 160 mM reaction additive 1 and 10 µl of 2 M reaction additive 2 (All components from the Click-iT Protein Enrichment Kit) were combined and added to each sample. Samples were then incubated at 40°C for 2 h while shaking, leading to the covalent capture of AHA-labeled proteins to the alkyne-activated resin. The beads were washed with water two times and resuspended in 100 µl water. In order to reduce and alkylate proteins bound to the beads, 2 ml SDS-wash buffer (Click-iT Protein Enrichment Kit) supplemented with 1 mM TCEP and 0.25 mM Chloroacetamide were added to each sample. Samples were then incubated at 70°C for 15 min while shaking at 1,800 rpm in an orbital shaker, followed by 15 min incubation at room temperature, also shaking at 1,800 rpm. Resins were transferred into a spin column (Click-iT Protein Enrichment Kit) and stringently washed with 2 ml water, 5x 2 ml SDS-wash buffer (Click-iT Protein Enrichment Kit), 2 ml water, 5x 2 ml guanidine wash buffer (100 mM Tris-HCL pH 8, 6 M Guanidine-HCl) and 5x 2 ml 20% (v/v) acetonitrile (ACN). The resin was resuspended in 2 ml digestion buffer (100 mM Tris-HCl pH8, 5% (v/v) ACN, 2 mM CaCl_2_) and pelleted by centrifugation. 1.8 ml of digestion buffer was removed, leaving the beads in 200 µl, and 10 µl of 0.1 µg/µl Trypsin/LysC Mix (Promega #V5073) was added. Proteins were digested at 37°C overnight while shaking at 1,000 rpm in an orbital shaker. To retrieve the tryptic peptides, the resin was pelleted and the supernatant transferred to a fresh tube. The resin was resuspended in 600 µl water, vortexed, pelleted and the supernatant added to the previous tube (=800 µl total). Samples were acidified by adding formic acid (FA) to 1% (v/v) and cleaned up via reversed-phase solid phase extraction with an Oasis PRiME HKB µElution Plate (Waters #186008052) following the instructions. Eluates were dried in a centrifugal evaporator and resuspended in 0.1% (v/v) FA. If peptide concentrations were measured, the Pierce^TM^ Quantitative Colorimetric Peptide Assay (Invitrogen #C10416) was used. Peptides were then fractionated via high-pH reversed-phase fractionation.

### High-pH reversed-phase fractionation

Samples were fractionated using an Infinity 1260 LC system (Agilent) with a Gemini C_18_ 3 µm-particle-size column (1 × 100 mm, Phenomonex #00D-4439-A0). An alkaline water/ACN gradient with a flow rate of 0.1 ml/min was used for elution. Solvent A contained 20 mM ammonium formate in H_2_O at pH 10 and solvent B was 100% ACN. The following gradient profile was used: 0-2 min 0% B, 2-60 min linear gradient to 65% B, 60-62 min linear gradient to 85% B, 62-67 min 85% B, 67-85 min 100% B. Eluting samples were either collected in 60 fractions and concatenated to 12 fractions or collected in 40 fractions and concatenated to 8 fractions. Samples were dried in a centrifugal evaporator and resuspended in 1% (v/v) FA. All fractions were cleaned up via reversed-phase solid phase extraction with an Oasis PRiME HKB µElution Plate (Waters #186008052) following the instructions. Eluates were dried in a centrifugal evaporator and resuspended in 0.1% (v/v) FA, followed by data dependent acquisition (DDA)-MS.

### DDA-MS for degradome and translatome analysis

An Easy-nLC1200 system (Thermo Scientific) connected to an Orbitrap Fusion Tribrid MS (Thermo Scientific) was used for DDA proteomics analysis. Peptides were loaded onto an Acclaim Pepmap C_18_ 5 µm-particle-size trap column (0.1 × 20 mm, Thermo Scientific #164564-CMD) and separated over a nanoEase M/Z Peptide BEH C_18_ 1.7 µm-particle-size analytical column (0.075 × 250 mm, Waters #186008795) which was heated to 55°C. Peptides were resolved with a water/ACN gradient consisting of solvent A (0.1% (v/v) FA in H_2_O) and solvent B (0.1% (v/v) FA and 80% (v/v) ACN in H_2_O). The following gradient profile was used with a flow rate of 300 nl/min: 0-2.8 min 3.7% B, 2.8-50 min linear gradient to 33.9% B, 50-58 min linear gradient to 52.4% B, 58-58.1 min linear gradient to 83.2% B, 58.1-74 min 83.2% B, 74-75 min linear gradient to 3.7% B, 75-105 min 3.7% B. Eluted ions were injected by electrospray ionization (ESI) and analyzed by MS/MS. The ion transfer tube temperature was set to 320°C and MS1 was performed at a resolution of 60,000 within the range of 375-1500 m/z. The AGC target was set to 1×10^6^ and the maximum injection time to 50 ms. The most abundant ions with a charge between 2-4 were selected from a 3 s cycle time window. Dynamic exclusion was set to 20 s with a mass tolerance of +-10 ppm. MS2 occurred with a higher-energy collisional dissociation (HCD) of 33% and an isolation window of 1.6 m/z. The AGC target was set to 1×10^4^ and the maximum injection time to 50 ms.

### Peptide search for degradome and translatome analysis

Proteins and peptides were inferred from raw spectral files via MaxQuant v1.6.0.16 (Cox & Mann, 2008) using a revised UniProt human proteome database (TaxID: 9609, Proteome ID: UP000005640) and the integrated default andromeda list of contaminants. SILAC specifications, the match-between-runs option and intensity-based absolute quantification (iBAQ, Schwanhäusser et al., 2011) were activated in all searches. The remaining settings were used at default.

### Sample preparation for whole-proteome analysis

Unlabeled Jurkat cells were treated for 8 hours either with 100 nM CC-885 or DMSO, harvested by centrifugation and washed with ice-cold PBS. Cell pellets were lysed in 0.1% RapiGest buffer (0.1% RapiGest in 100 mM Ammonium bicarbonate, 40 mM TCEP and 40 mM Chloroacetamide) and sonicated using a probe sonicator. Protein concentrations were determined from cleared lysates with either the Pierce^TM^ BCA Protein Assay Kit (Thermo Scientific #23225) or the Pierce^TM^ Rapid Gold BCA Protein Assay Kit (Thermo Scientific #A53225PR). 100 µg of protein was digested overnight at 37°C using Trypsin/LysC. On the next day, digestion was stopped by adding TFA to a final concentration of 1% (v/v). Samples were then incubated at 37°C for 30 minutes and centrifuged at 16000x g and 4°C for additional 30 minutes. Cleared supernatants were collected and analyzed by data independent acquisition (DIA)-MS.

### DIA-MS for whole-proteome analysis

An Easy-nLC1200 system (Thermo Scientific) connected to a Q Exactive HF MS (Thermo Scientific) was used for DIA proteomic analysis. Peptides were loaded onto an Acclaim Pepmap C_18_ 5 µm-particle-size trap column (0.1 × 20 mm, Thermo Scientific #164564-CMD) and separated over a nanoEase M/Z Peptide BEH C_18_ 1.7 µm-particle-size analytical column (0.075 × 250 mm, Waters #186008795) which was heated to 55°C. Peptides were resolved with a water/ACN gradient consisting of solvent A (0.1% (v/v) FA in H_2_O) and solvent B (0.1% (v/v) FA and 80% (v/v) ACN in H_2_O). The following gradient profile was used with a flow rate of 300 nl/min: 0-3 min 2% B, 3-153 min linear gradient to 25% B, 153-183 min linear gradient to 40% B, 183-184 min linear gradient to 95% B, 184-189 min 95% B, 189-190 min linear gradient to 2% B, 190-210 min 2% B. Eluted ions were injected by electrospray ionization (ESI) and analyzed by MS/MS. The ion transfer tube temperature was set to 320°C and MS1 was performed at a resolution of 60,000 within the range of 400-1200 m/z. The automatic gain control (AGC) target was set to 3×10^6^ and the maximum injection time to 20 ms. Data-independent MS/MS spectra were acquired at an Orbitrap resolution of 30,000, an AGC target of 3×10^6^ and an automated maximum injection time using 34 non-overlapping isolation windows of 24.3 m/z. The normalized collision energy was set to 27 and the fixed first mass to 200 m/z.

### Peptide search for whole-proteome analysis

Proteins and peptides were inferred from raw spectral files via Spectronaut v15.1.210713 (Bruderer et al., 2015) in directDIA mode using a revised UniProt human proteome database (TaxID: 9609, Proteome ID: UP00000564). The following search settings were changed in comparison to default settings: Proteins were digested with Trypsin and LysC.

### Data Filtering and limma analysis

Proteins and peptides that were marked by MaxQuant as *contaminants*, *reverse* or *only identified by site* were excluded from all datasets. The limma R package v.3.44.3 was used to analyze differential protein expression (Ritchie et al., 2015).

### Luminescence assay

For NanoLuc measurements, 10000 cells in 30 µl media were seeded into each well of a black 384-well plate (Corning #3764). 1:100 Endurazine Luciferase live cell substrate (Promega) were added to each well, followed by a 60 min incubation. At time point 0, 10 µl of pre-made compound solution in media was added to each well and luciferase measurements were performed on an EnVision plate reader (PerkinElmer).

### Data availability

The mass spectrometry proteomics data have been deposited to the ProteomeXchange Consortium via the PRIDE (Perez-Riverol et al., 2022) partner repository with the dataset identifier PXD048931.

## Acknowledgements

This work was funded in part by the Deutsche Krebshilfe (project 70114190) (to J.K.). CeMM and the Winter Lab are supported by the Austrian Academy of Sciences. The Winter lab is supported by funding from the European Research Council (ERC) under the European Union’s Horizon 2020 research and innovation program (grant agreement 851478), as well as by funding from the Austrian Science Fund (FWF, projects P32125 and P7909). Figure 1a and Supplementary Figure S2 were created with biorender.com.

## Author contributions (CRediT)

Conceptualization: M.J., J.K.; Formal analysis: M.J., A.S., L.M.W; Funding acquisition: J.K., G.E.W.; Investigation: M.J., A.S.; Methodology: M.J., J.K.; Resources: J.C., A.N.; Supervision: J.K., G.E.W. Writing-original draft: M.J. Writing-review and editing: J.K., G.E.W., A.S., L.M.W.

---

**Figure S1:**
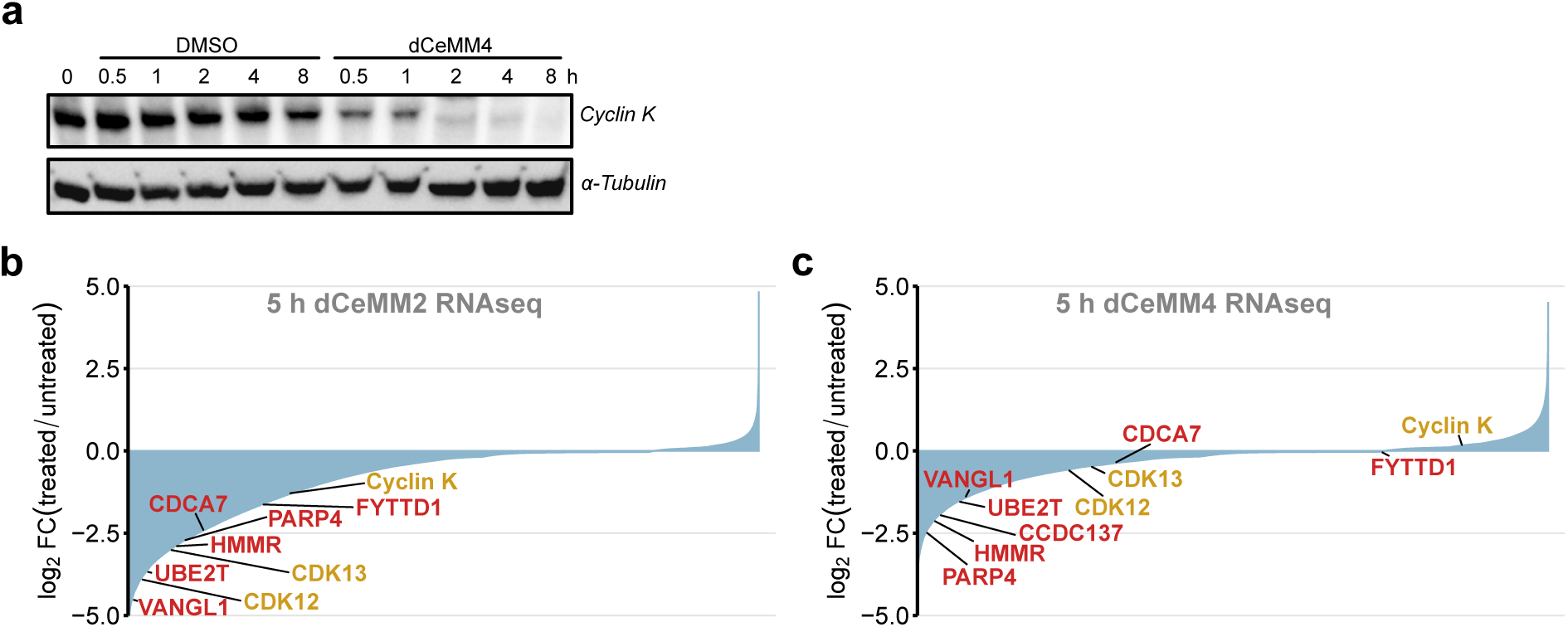
**(a)** Degradation of Cyclin K in response to dCeMM4 determined by immunoblot analysis. KBM7 cells were either treated with 2.5 µM dCeMM2 or with DMSO for the indicated time intervals. Upper panel: anti-Cyclin K antibody. Lower panel: anti-α-Tubulin (loading control). **(b)** RNAseq analysis from Mayor-Ruiz et al., 2020 showing the transcriptome response to 5 h dCeMM2 (2.5 µM) or **(c)** to dCeMM4 (3.5 µM) treatment. Highlighted in yellow are the three known targets of dCeMM2 and dCeMM4: Cyclin K, CDK12 and CDK13. Highlighted in red are the proteins that exhibit both a more extreme fold-change and a better p-value than Cyclin K, CDK12 and CDK13 in the corresponding whole proteome dataset, and that were identified using degradome proteomics (see Figure 2a and b).

**Figure S2:**
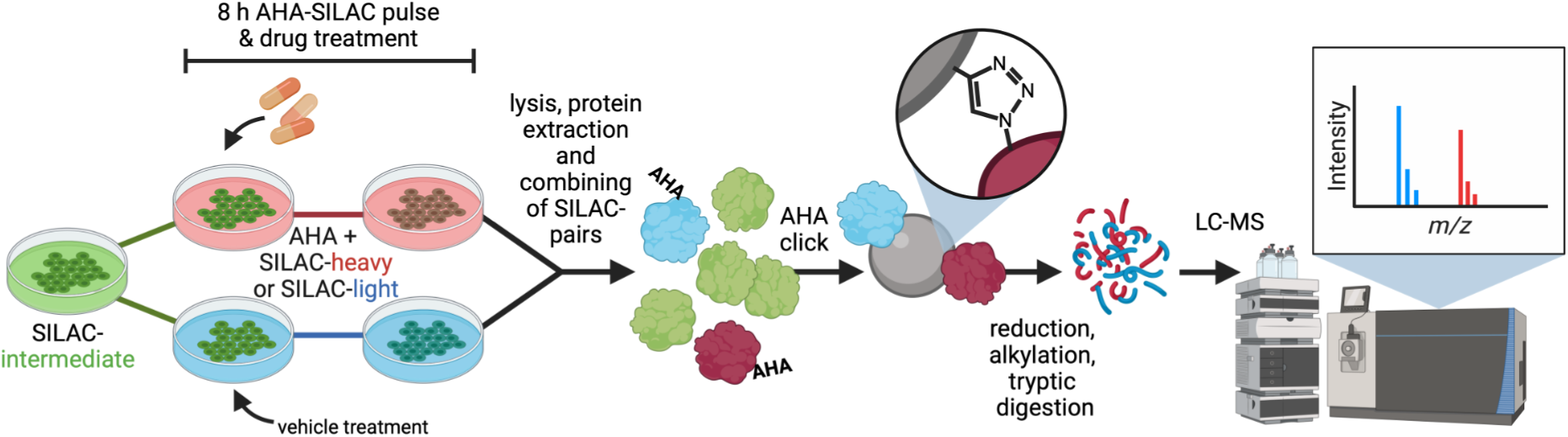
Scheme for nascent proteome analysis. Fully SILAC-intermediate labeled cells are incubated in both AHA and SILAC-heavy or SILAC-light amino acids containing media. The heavy-channel is treated with the drug of interest, while the light-channel is mock-treated. After 8 h, cells are lysed, proteins extracted, and the corresponding SILAC-channels combined. The pulse-labelled proteins are enriched for AHA via click-chemistry, reduced, alkylated, and digested by trypsin, followed by MS analysis. Heavy-over-light SILAC-ratios are used to identify drug-induced changes in the newly synthesized proteome.

